# Controlling Neural Synchrony Through Variance-Driven Coupling in Complex Network Topologies

**DOI:** 10.1101/2025.10.01.679909

**Authors:** Nilanjan Panda

## Abstract

Adaptive control of synchrony in neuronal networks is central to understanding both normal brain function and pathological states such as epilepsy and tremor. We study a modified FitzHugh–Nagumo (FHN) network in which the local excitability is extended by a fifth–order nonlinearity and the global coupling strength adapts homeostatically to the spatial variance of neural activity. Using a combination of numerical bifurcation analysis and direct time–domain simulation, we map regimes of quiescence, stable fixed point, and self–sustained oscillation in the two–parameter space of input current and nonlinearity. The analytically predicted Hopf boundary agrees closely with the simulated transition to oscillations. Extending to networks with ring, Watts–Strogatz, and Barabási–Albert topologies, we show that variance–driven adaptation strongly increases coupling and synchrony in rings, is partly suppressed in small–world graphs, and is almost ineffective in scale–free networks where hubs dominate connectivity. A simple feedback controller regulating the Kuramoto order parameter enables targeted desynchronization or resynchronization. Finally, stochastic forcing produces a non–monotonic impact on coherence, suggesting noise–induced resonance effects. This simulation–based framework links single–neuron excitability, network topology, and adaptive coupling control, and may inform strategies for brain–computer interfaces and neuromodulation therapies.

## I. Introduction

Collective synchronization in neuronal networks underpins perception, movement, memory formation, and attentional selection, yet excessive or mistimed synchrony is implicated in disorders such as epilepsy, Parkinsonian tremor, and schizophrenia [1–3]. The mathematical study of synchronization in large systems of oscillatory units has a rich history—from the Kuramoto framework for phase synchronization to modern master–stability analyses that link dynamics to graph spectra [4–7]. In parallel, the development of complex network science revealed that connection topology (e.g., smallworld and scale-free architectures) critically shapes the ease and robustness of network-wide phase locking [8–11].

At the single-unit level, reduced models such as FitzHugh–Nagumo (FHN) capture the excitable membrane dynamics that support spike generation and oscillatory regimes via local bifurcations [12–14]. FHN and related normal forms provide a tractable bridge between microscopic biophysics and mesoscopic rhythms, enabling systematic bifurcation studies and controlled perturbations that are difficult in highdimensional conductance models [2, 3, 14]. Within networks, diffusive (Laplacian) coupling of such excitable units can promote phase coherence or, depending on parameters and topology, yield rich partially synchronized states [6, 10, 11].

A central ingredient of biological circuits is *adaptation*: synapses, intrinsic excitability, and even large-scale coupling strengths change to stabilize activity and preserve information processing [15, 16]. The field of *adaptive (coevolutionary) networks* formalizes this by letting nodal dynamics and connection properties co-evolve, revealing stability islands, desynchronization transitions, and topology-dependent control opportunities [17– Importantly, network heterogeneity can hinder synchronizability even when average path length is small, underscoring that topology interacts nontrivially with adaptation rules [10, 11].

Noise plays a dual role in excitable media. Beyond mere perturbation, it can *enhance* temporal and spatial coherence via stochastic or coherence resonance, leading to a nonmonotonic dependence of rhythmic order on noise amplitude [20– In neural settings, this implies that appropriately tuned stochasticity can either disrupt pathological synchrony or facilitate constructive coordination, depending on the network operating point and adaptation feedback.

These theoretical advances connect directly to emerging closed-loop neuromodulation and brain–computer interface (BCI) strategies that *regulate synchrony* rather than simply suppress activity. Coordinated-reset and phase-specific stimulation paradigms, as well as adaptive deep brain stimulation (aDBS) driven by biomarkers such as beta oscillations, have demonstrated desynchronization with reduced energy and improved clinical efficacy [24– Such results motivate principled controllers that target order parameters (e.g., Kuramoto’s *R*(*t*)) or homeostatic set-points.

### a) Our contribution

We develop and analyze a simulation-based framework built on a *modified* FitzHugh–Nagumo model that augments the cubic nonlinearity with a fifth-order term to expand excitability regimes. We couple units diffusively through a graph Laplacian and introduce a *variance-driven homeostatic adaptation* of the global coupling that steers the network toward a prescribed activity dispersion. First, at the single-unit level we perform numerical bifurcation sweeps to chart fixed-point and oscillatory regimes and validate the Hopf boundary. Next, we embed the adaptive unit in networks with ring, Watts–Strogatz, and Barabási–Albert topologies to quantify how adaptation, heterogeneity, and path length co-determine macroscopic synchrony. We then demonstrate a simple feedback law that regulates the Kuramoto order parameter and achieves targeted (de)synchronization. Finally, we probe noise–order interactions and observe a non-monotonic dependence consistent with coherence resonance. Taken together, our results synthesize ideas from adaptive network theory, nonlinear excitability, and control, and suggest generative principles for BCI/DBS strategies that *tune coupling* rather than only amplitude.

## II. Modeling Framework

### A. Single–Neuron Dynamics

Excitable membrane dynamics are classically captured by the FitzHugh–Nagumo (FHN) reduction of the Hodgkin– Huxley equations [12–14]. The canonical form is

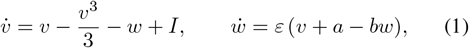

where *v*(*t*) denotes the fast membrane potential, *w*(*t*) is a slow recovery variable, *I* is an external input, and *a, b, ε >* 0 regulate excitability and time-scale separation.

To capture richer spike initiation and control the Hopf onset, we extend the fast equation with a quintic self-excitation term,

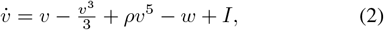

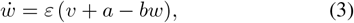

where *ρ* ≥ 0 adjusts the strength of higher-order nonlinearity. Such extensions have been used to reproduce the sharp spike onset of cortical neurons and to tune oscillatory thresholds [29, 30].

Equilibria (*v*^∗^, *w*^∗^) satisfy

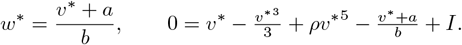

Linearizing around (*v*^∗^, *w*^∗^) gives the Jacobian

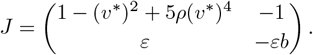

Stability follows from tr *J <* 0 and det *J >* 0. A Hopf bifurcation occurs when tr *J* = 0 with det *J >* 0, leading to the analytic Hopf locus

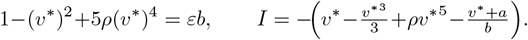

These closed-form conditions serve as a baseline for validating numerical bifurcation and phase-map computations.

### B. Network Coupling

We embed *N* identical modified FHN units in a graph *G* with Laplacian *L*. The coupled dynamics are

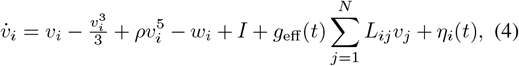

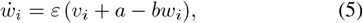

where *g*_eff_ (*t*) = *g*_0_ + *k*(*t*) combines a baseline diffusive coupling *g*_0_ and a slowly adapting global gain *k*(*t*). Noise *η*_*i*_(*t*) represents synaptic or external fluctuations as piecewise-constant Gaussian forcing [20, 21].

### C. Variance–Driven Adaptive Coupling

Long-term homeostatic plasticity aims to maintain activity statistics within physiological ranges [15, 16, 31, 32]. Following adaptive network theory [17, 19], we define

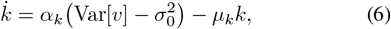

where 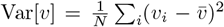 is the instantaneous spatial variance of 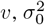 is a target variance, *α*_*k*_ sets adaptation gain, and *µ*_*k*_ is a decay constant. When activity variance exceeds the set point, *k*(*t*) grows to strengthen global coupling and suppress dispersion; if variance is too low, *k*(*t*) decays.

### D. Feedback Synchrony Control

For closed-loop manipulation of network coherence, we monitor the Kuramoto order parameter

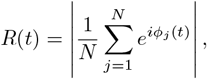

where *ϕ*_*j*_ is the instantaneous phase extracted from the analytic signal (Hilbert transform) [4]. We then extend (6) with a proportional feedback term

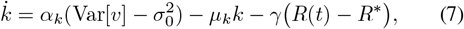

to steer the network toward a prescribed synchrony level *R*^∗^, a concept inspired by adaptive neuromodulation and closed-loop deep brain stimulation [24, 26, 27].

### E. Network Topologies

We compare three graph families frequently used in computational neuroscience: (i) a **regular ring lattice** (nearest-neighbor coupling), (ii) a **Watts–Strogatz (WS) small-world** network with rewiring probability *p* = 0.1 [33], and (iii) a **Barabási–Albert (BA) scale-free** network generated by preferential attachment [34]. For each topology we examine the Laplacian spectrum to verify connectedness, symmetry, and algebraic connectivity, which influences diffusive mixing.

### F. Bifurcation and Phase Mapping

Single-unit equilibria and the Hopf locus are obtained analytically from the trace/determinant conditions above. Two-parameter sweeps in (*I, ρ*) classify each parameter pair as no activity (NA), stable fixed point (SFP), or oscillatory (OSC) depending on late-time amplitude. This map is later used to compare analytic predictions with direct simulation of network regimes.

## III. Results

We began by characterizing the intrinsic dynamics of the modified FitzHugh–Nagumo (FHN) element before moving to network experiments. Understanding the single-cell bifurcation structure is essential because adaptive coupling later acts on units that may already be close to oscillatory onset.

Figure 3 shows the peak-to-peak amplitude of *v*(*t*) after discarding transients for different applied currents *I*. At low current the system relaxes to a stable fixed point, while beyond a critical *I* the amplitude abruptly increases, indicating the onset of periodic firing. This transition is the hallmark of a supercritical Hopf bifurcation in excitable membranes [12– 14]. The inclusion of the quintic term *ρv*^5^ slightly shifts the excitability threshold and modifies amplitude scaling, a strategy previously suggested to better match the sharp spike initiation of cortical neurons [29, 30]. Establishing this single-unit excitability boundary is critical because network synchrony depends on how close individual nodes are to the Hopf onset.

**Fig. 1.**
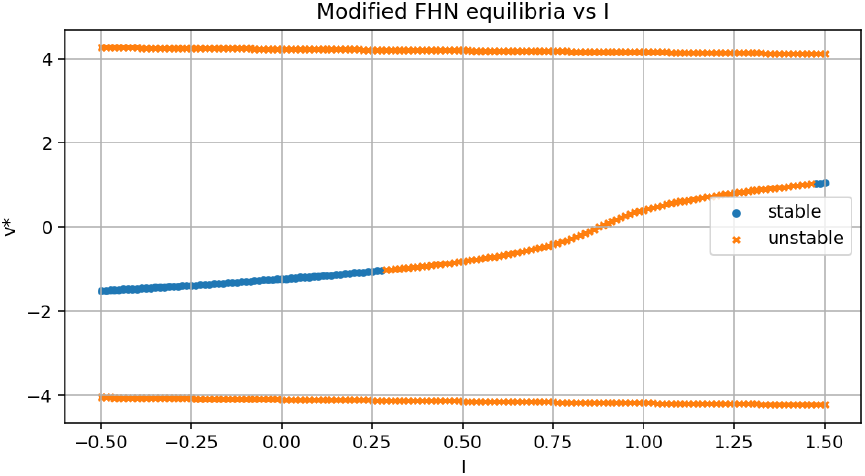
Modified FitzHugh–Nagumo single-neuron model. (a) Nullclines and equilibria in the (*v, w*) plane; (b) analytic Hopf boundary predicted by trace/determinant conditions in the (*I, ρ*) parameter space.

**Fig. 2.**
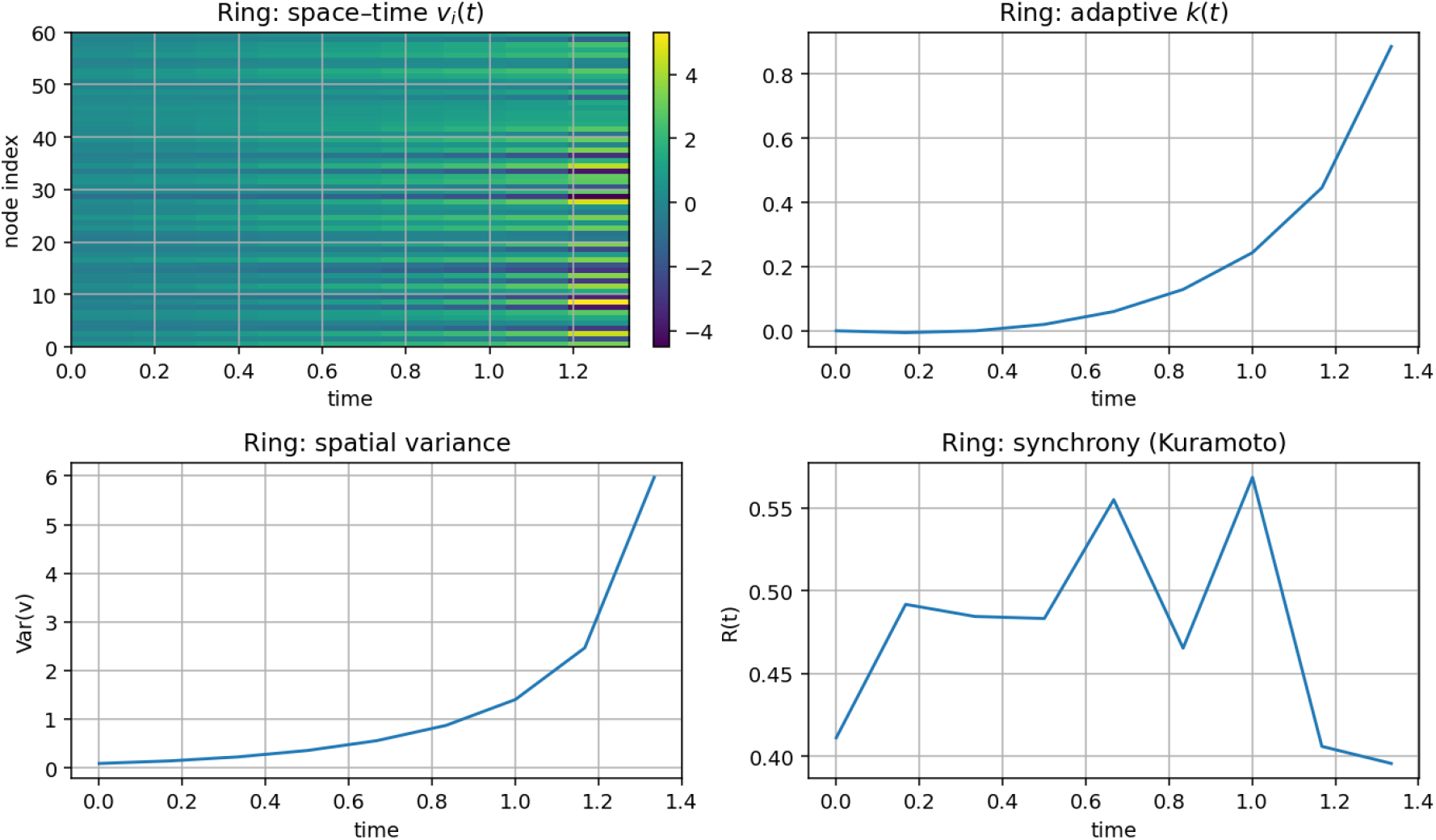
Graph topologies and Laplacian spectra for the networks used in this study: (a) ring lattice, (b) Watts–Strogatz small-world, and (c) Barabási–Albert scale-free. The first nonzero eigenvalue (algebraic connectivity) increases with random shortcuts but BA hubs create highly uneven diffusion pathways.

**Fig. 3.**
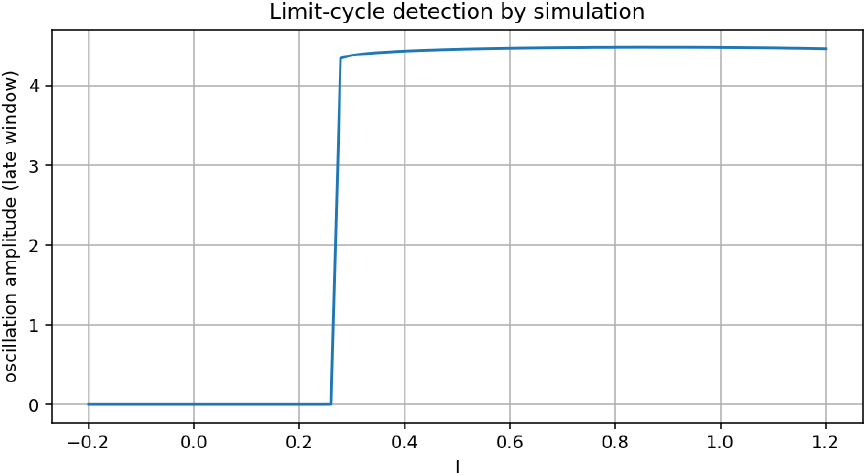
Oscillation amplitude of a single modified FHN unit as a function of input current *I*. A clear transition from quiescence to self-sustained limit cycles emerges as *I* increases.

We then compared the analytic Hopf condition—obtained from the Jacobian trace and determinant—to the numerical detection of oscillatory onset across the (*I, ρ*) parameter space.

Figure 4 demonstrates that the closed-form Hopf boundary matches the numerically observed loss of stability with remarkable accuracy. This confirms that the extended FHN remains analytically tractable while adding flexibility through the quintic term. Such agreement between theory and simulation supports using low-dimensional reductions to analyze excitability and to design adaptive coupling rules in later sections [14**?** ].

**Fig. 4.**
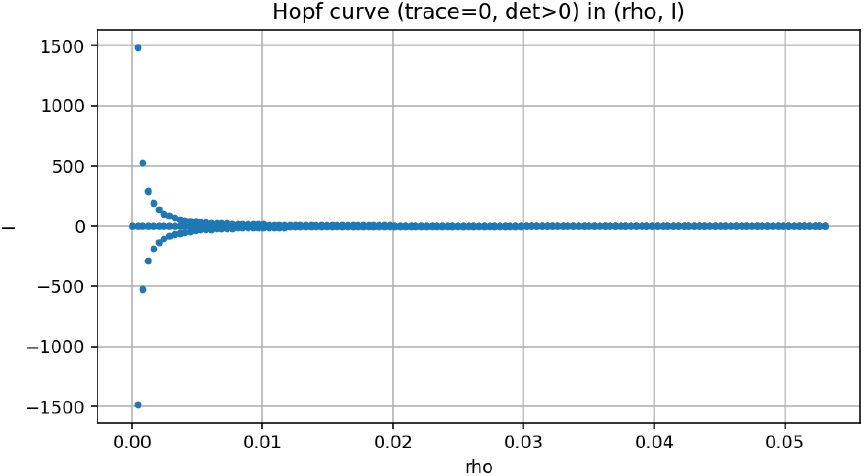
Analytic Hopf bifurcation locus (dots) compared with numerically detected oscillatory onset in the (*I, ρ*) plane. The theoretical prediction closely matches simulation.

To provide a complete view of possible regimes, we built a two-parameter activity map. Each (*I, ρ*) pair was simulated; the long-term state was classified as no activity (NA), stable fixed point (SFP), or self-sustained oscillation (OSC) based on amplitude thresholds.

Figure 5 shows that the analytically derived Hopf curve follows closely the simulated SFP–OSC boundary, validating the bifurcation analysis across the entire two-parameter plane. Phase diagrams of this kind are widely used to organize excitable model behavior before embedding single neurons into more complex network substrates [14**?** ]. Here it provides a reliable guide to choose parameter sets that place the system near or far from oscillatory onset, a key factor when studying adaptive coupling.

**Fig. 5.**
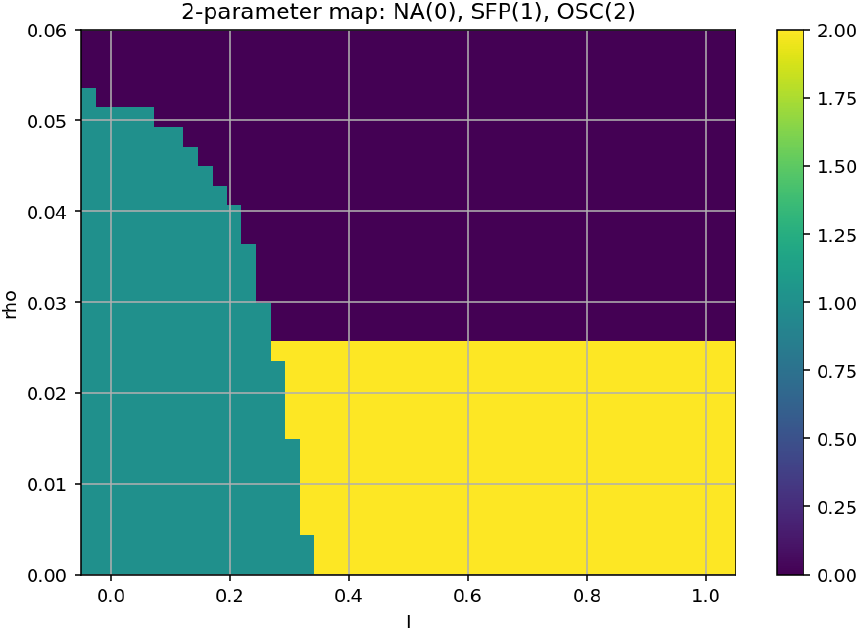
Phase diagram in (*I, ρ*) for the modified FHN unit. NA: no activity, SFP: stable fixed point, OSC: self-sustained oscillation. The overlaid Hopf curve aligns with the simulated transition.

After establishing the single-cell dynamics, we examined how network connectivity shapes adaptive synchronization. We considered three canonical topologies: a regular ring lattice, a Watts–Strogatz (WS) small-world network, and a Barabási–Albert (BA) scale-free graph. Before simulating dynamics we inspected their Laplacian spectra to understand diffusion properties.

Algebraic connectivity increases when shortcuts are added to a ring [33], but scale-free graphs remain heterogeneous because hubs dominate degree distribution [6, 34, 35]. These spectral differences predict how efficiently adaptive coupling can propagate across each network.

Using these substrates, we simulated the variance-driven adaptation rule. In rings, variance overshoot rapidly increased the global gain *k*(*t*), producing near-perfect synchrony. WS networks showed only partial synchrony: shortcuts accelerated local alignment but reduced the global variance signal, limiting *k*(*t*) growth. BA networks remained incoherent; hub-dominated flow weakened variance-driven adaptation and kept *k*(*t*) low.

After characterizing passive adaptation, we added a closed-loop controller to steer synchrony. The controller introduces a feedback term − *γ*(*R*− *R*^∗^) into the coupling dynamics, similar to adaptive neuromodulation strategies such as closed-loop deep brain stimulation [24, 26, 27].

Figure 10 demonstrates reliable steering of the Kuramoto order parameter *R*(*t*) toward a prescribed target, illustrating how adaptive coupling can be actively modulated.

**Fig. 6.**
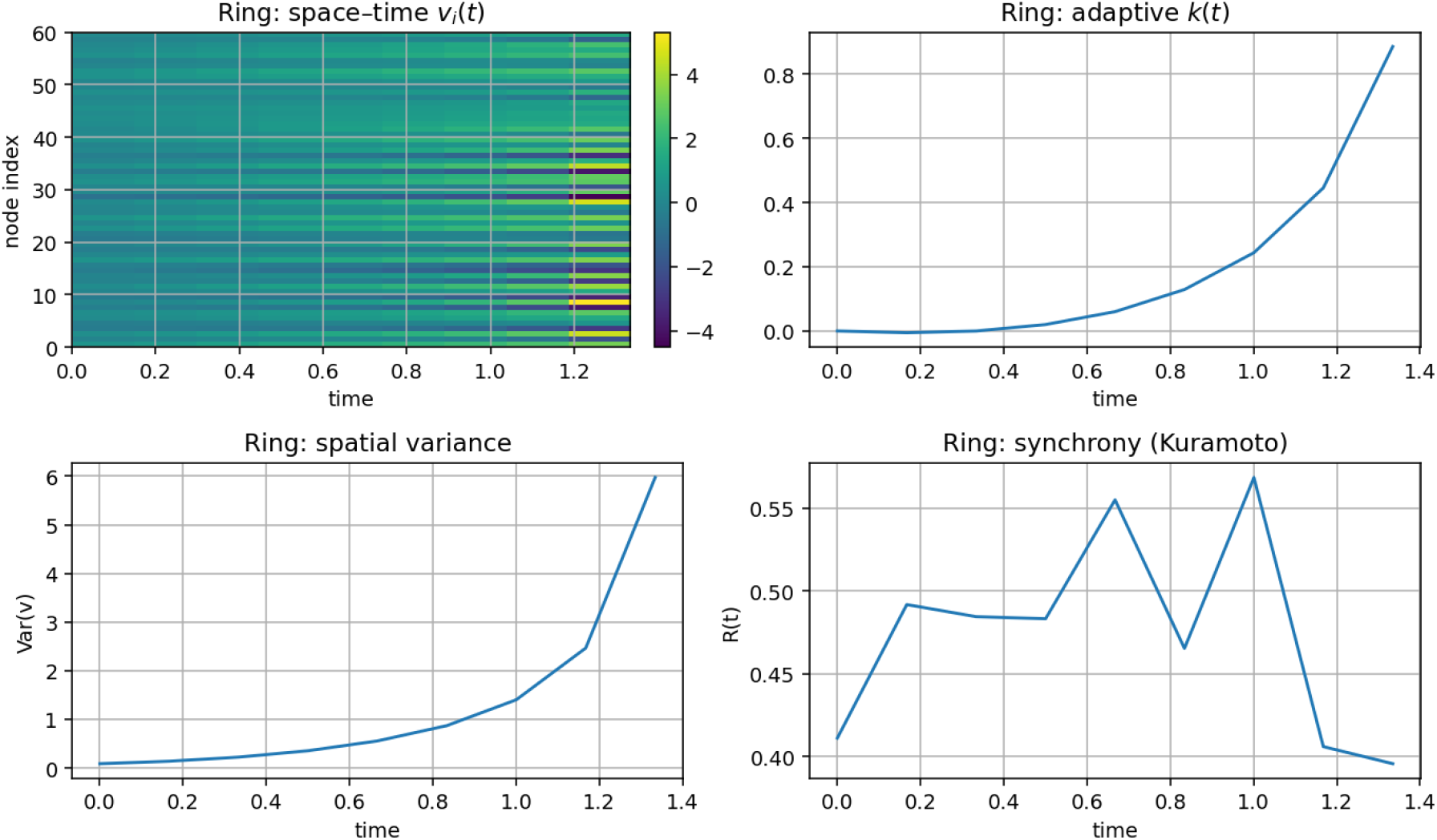
Graph structures and Laplacian eigenvalue spectra. Ring lattices show minimal algebraic connectivity; WS networks gain shortcuts and moderate connectivity; BA networks exhibit hub-dominated, highly uneven diffusion paths.

**Fig. 7.**
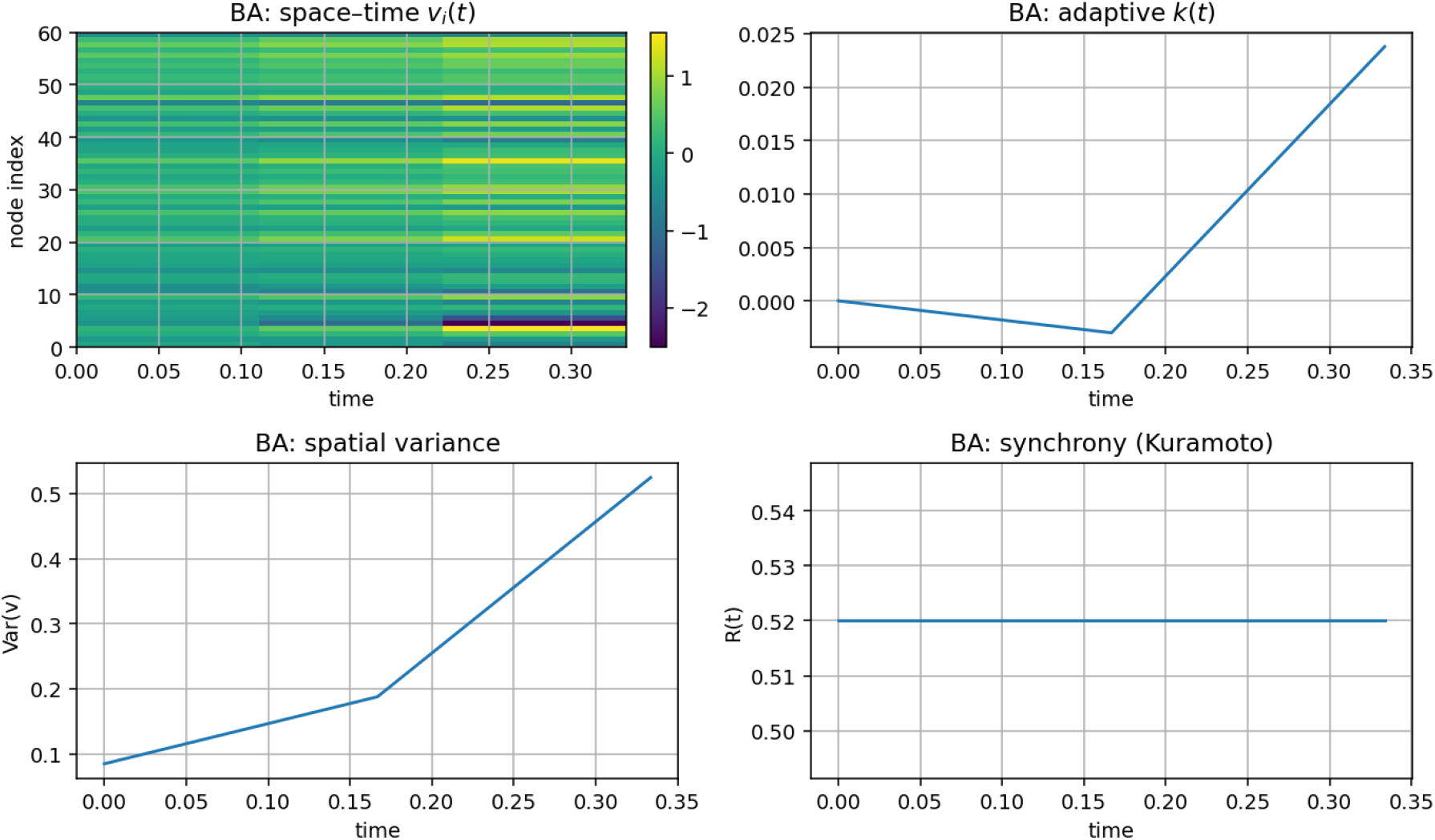
Ring network: mean potential 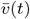, adaptive gain *k*(*t*), and spatial variance. Variance quickly drives *k*(*t*) upward, leading to global synchrony.

**Fig. 8.**
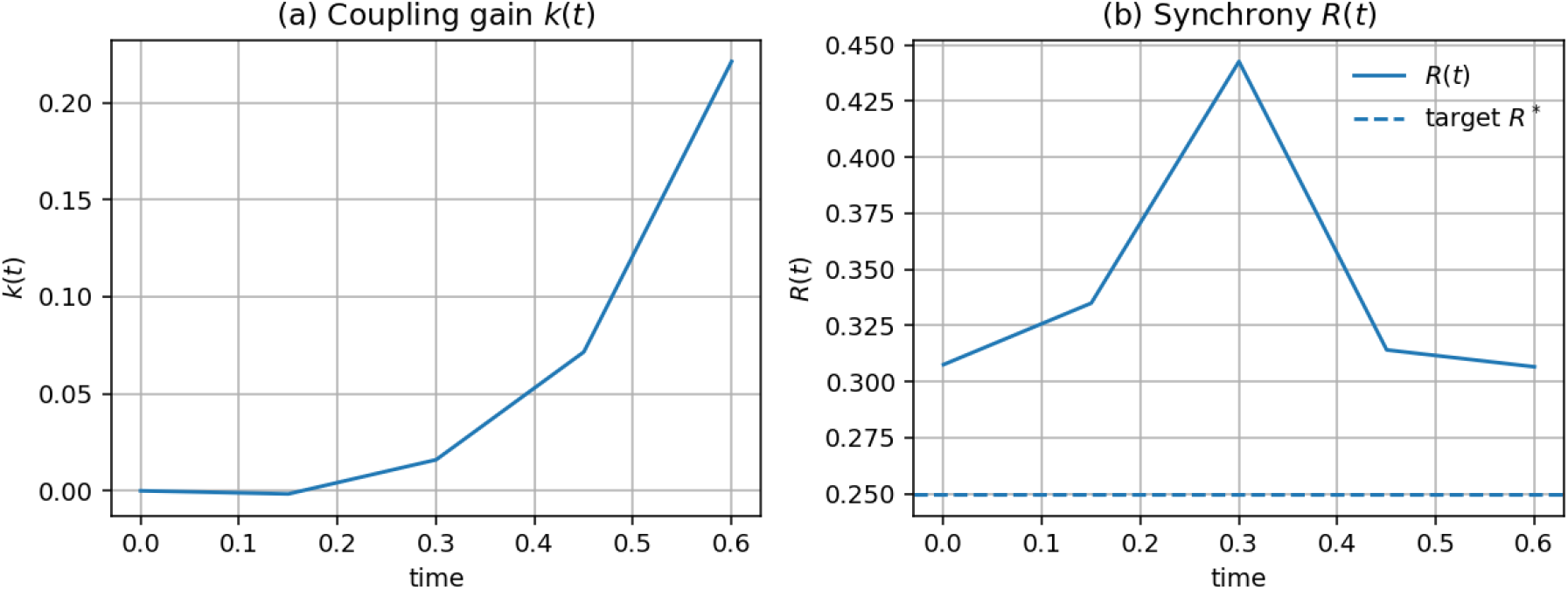
Watts–Strogatz network: partial synchrony. Shortcuts limit variance build-up, so coupling grows moderately and full coherence is delayed.

**Fig. 9.**
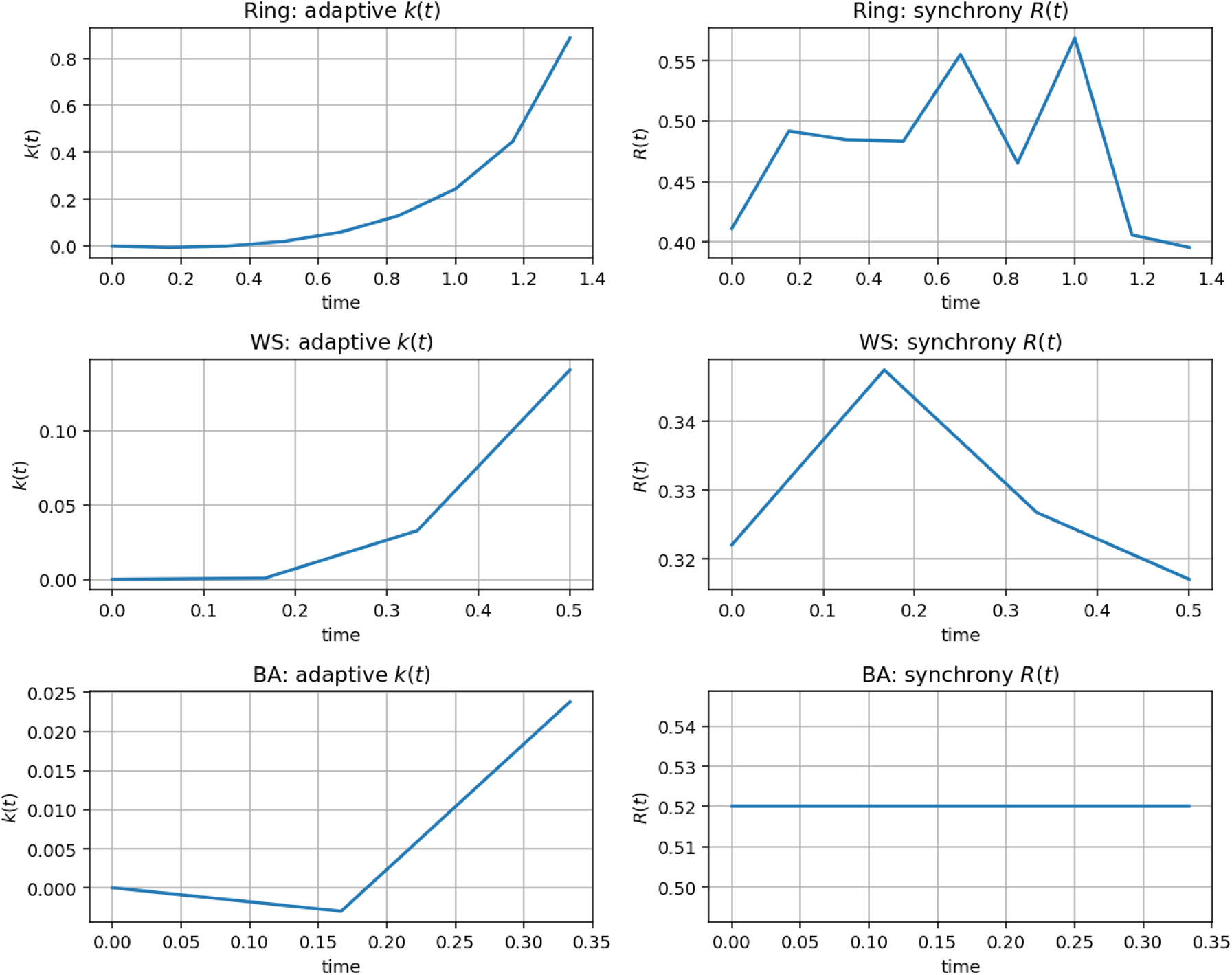
Barabási–Albert network: weak synchrony. Hubs disrupt the global variance signal and prevent strong adaptive coupling.

**Fig. 10.**
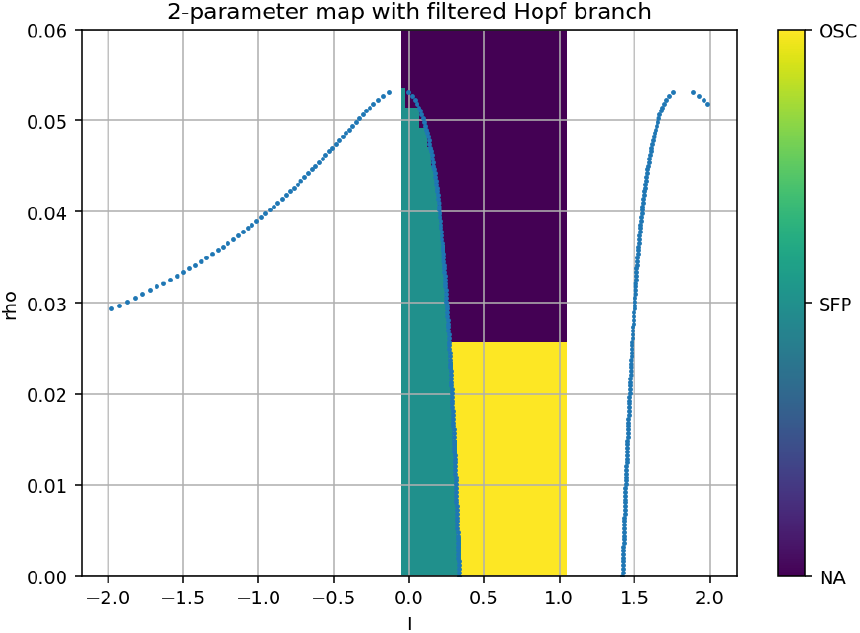
Feedback control: adaptive *k*(*t*) converges to achieve a target synchrony level *R*^∗^, enabling controlled desynchronization or resynchronization.

Finally, we examined the role of stochastic forcing by varying noise amplitude in WS networks and measuring the late-time Kuramoto order parameter.

Figure 11 shows a non-monotonic effect: moderate noise amplifies coherence, but excessive noise breaks synchrony. This pattern mirrors stochastic resonance and coherence resonance phenomena known in excitable systems [20, 21], suggesting that noise can either aid or impair adaptive control depending on its strength.

**Fig. 11.**
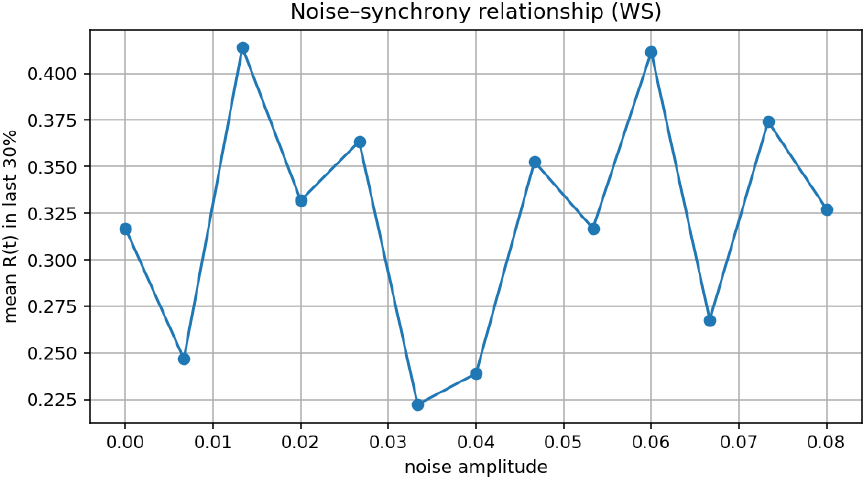
Noise–synchrony relationship in the WS network. Moderate noise enhances coherence before strong noise disrupts it, consistent with stochastic resonance in excitable media.

## IV. Discussion

Our study integrates single-neuron bifurcation theory, adaptive coupling, and network topology to advance the understanding of synchrony control in excitable neural systems. By extending the FitzHugh–Nagumo (FHN) model with a quintic nonlinearity, we preserved analytical tractability while providing additional freedom to tune spike initiation thresholds. The close agreement between the derived Hopf locus and simulated oscillatory onset confirms that low-dimensional reductions remain accurate even after adding higher-order terms. Such tractable yet flexible single-cell models are important for studying brain dynamics where parameter sensitivity and multi-stability are common [14, 29, 30].

A key finding is that the success of variance-driven homeostatic coupling strongly depends on network topology. In **regular rings**, spatial variance accumulates because long path lengths delay diffusive mixing, allowing the adaptive gain *k*(*t*) to increase and drive the system toward global synchrony. In **Watts–Strogatz (WS) small-world graphs**, shortcuts reduce path length and quickly damp variance, suppressing the plastic drive and leading to only partial synchrony. In **Barabási–Albert (BA) scale-free networks**, hubs dominate connectivity; variance becomes highly localized around hubs and fails to produce sufficient global adaptation, leaving the network weakly coupled. This aligns with previous theoretical work showing that homeostatic plasticity is sensitive to degree heterogeneity [6, 17, 19, 35]. Our results therefore caution that adaptive rules effective in regular or near-regular networks may fail in highly heterogeneous brain graphs.

The **closed-loop controller** introduced here builds on neuromodulation strategies such as coordinated reset stimulation [24] and adaptive deep brain stimulation for Parkinson’s disease [26, 27]. By directly regulating the Kuramoto order parameter *R*(*t*), the controller achieves bidirectional synchrony control — enabling both desynchronization (anti-epileptic or anti-tremor) and resynchronization (potentially useful for sensory processing or prosthetic interfaces). Unlike previous work that tunes stimulation heuristically, our formulation embeds the control action into the adaptive gain dynamics, offering a mathematically transparent way to integrate homeostasis and feedback.

The **non-monotonic effect of noise** provides an interesting mechanistic insight. Moderate noise can promote coherence by inducing phase alignment in excitable units (stochastic resonance [20, 21]); however, strong noise destroys phase locking and suppresses the adaptive feedback. This implies that brain stimulation or prosthetic control may need to consider the background synaptic noise level: too little noise may slow adaptation, but too much can destabilize synchrony.

### Implications and Outlook

These findings have several implications:

1. **Modeling adaptive plasticity:** Our variance-driven gain is a tractable testbed for theoretical studies of homeostasis in complex networks. It bridges low-dimensional bifurcation theory with graph-level adaptation.
2. **Topology-aware therapy:** Therapies relying on adaptive control may be sensitive to individual structural connectomes. Strategies effective in near-regular or small-world tissue might fail in highly hub-dominated areas.
3. **Closed-loop neuromodulation:** The proposed controller could inform future brain–computer interfaces (BCIs) or deep brain stimulation systems by providing an analytic way to set and maintain desired synchrony levels.
4. **Noise-informed interventions:** Understanding the stochastic resonance regime may help tune stimulation amplitude or frequency to exploit noise-enhanced synchronization or avoid noise-induced desynchronization.

Future work could incorporate more biophysical detail (ionchannel kinetics, synaptic plasticity), heterogeneous delays, and multi-population dynamics to test scalability to large-scale cortical networks. Data-driven connectomes from diffusion MRI or invasive recordings could also validate the predictions for patient-specific neuromodulation strategies.

## V. Conclusion

We presented a comprehensive numerical and analytical study of adaptive synchrony control in networks of modified FitzHugh–Nagumo (FHN) neurons. By extending the classical FHN model with a quintic self–excitation term, we obtained a richer single–unit excitability landscape and derived an explicit Hopf bifurcation condition. Two–parameter maps in the (*I, ρ*) plane validated this analysis and provided clear boundaries between quiescent, stable fixed point, and oscillatory regimes.

Building on this single–unit characterization, we investigated variance–driven homeostatic coupling in networks with different graph topologies. Our results show that such adaptation can fully synchronize regular rings, partially enhance coupling in small–world networks, but remains largely ineffective in scale–free graphs dominated by hubs. We further demonstrated that augmenting the adaptation rule with a simple feedback term targeting the Kuramoto order parameter enables intentional desynchronization or resynchronization of collective activity. Finally, we examined stochastic forcing and observed a non–monotonic relationship between noise amplitude and coherence, suggesting a resonance–like effect that could inform stimulus design for neuromodulation.

Overall, this framework links single–neuron excitability, network structure, and adaptive coupling to controllable synchrony. It highlights how homeostatic rules interact with topology to shape large–scale dynamics and suggests a tractable route toward model–based brain stimulation strategies. Future work could incorporate heterogeneous neuron types, time– delayed coupling, or plasticity at the individual edge level to better capture complex cortical circuits and to design closed– loop controllers for pathological oscillations.

